# Different computations over the same inputs produce selective behavior in algorithmic brain networks

**DOI:** 10.1101/2021.02.04.429372

**Authors:** Katarzyna Jaworska, Nicola J. van Rijsbergen, Robin A.A. Ince, Philippe G. Schyns

**Affiliations:** Institute of Neuroscience and Psychology, University of Glasgow, 62 Hillhead Street, G12 8QB, Glasgow; Department of Psychology, Edge Hill University, St Helens Road, L39 1QP, Ormskirk

## Abstract

A key challenge in systems neuroscience remains to understand where, when and now particularly *how* brain networks compute over sensory inputs to achieve behavior. We used XOR, OR and AND functions as behavioral tasks, because each requires a different computation over the same inputs to produce correct outputs. In each task, source-localized magnetoencephalographic activity progresses through four systems-level computations identified within individual participants (N = 10/task): (1) linear discrimination of each visual input, first contra-laterally in occipital cortex then (2) jointly in midline occipital cortex and right fusiform gyrus, followed by (3) nonlinear task-dependent input integration in temporal-parietal cortex and finally (4) behavioral response representation in post-central gyrus. Our results show how network algorithms differently compute over the same inputs to produce different behaviors.

**One sentence summary:** Four stages of task-specific computations over the same visual inputs achieve different behaviors in dynamic brain networks

---

Extensive studies revealed that the primate visual system comprises the ventral and dorsal pathways, with specific anatomical and functional hierarchical organization (*1, 2*). These pathways compute over the high-dimensional visual input, starting separately in each hemisphere with contra-lateral detection of simple, small features with small receptive fields, that are then hierarchically integrated into more complex, broader receptive field features (*3–5*), leading to the integrated face, object and scene features (*6–9*) that are compared with memory to produce behavior (*10–13*). This flow of information reverses when the same pathways predict the input topdown from memory (*14–16*).

There is broad agreement that such a bi-directional hierarchical architecture supports much of the information processing that subtends everyday face, object and scene recognition. However, despite considerable progress, we have yet to understand where, when and how specific algorithmic computations in the pathways dynamically represent and transform the visual input into integrated features to produce behavior (*17–19*)–and vice-versa, when reversing the flow in the hierarchy, to predict a cascade of features from complex to simpler ones. Further, it is unclear that such an algorithmic understanding can be achieved with current analytical approaches to neuroimaging, even in simple tasks (*20*). Here, we achieved such systems-level algorithmic understanding with MEG measurements, in the context of well-defined visual inputs and tasks.

We framed this broad problem using the classic logical functions XOR, AND and OR, in which different algorithms are required to produce correct responses from the same input stimuli (see these input-output relationships in Figures 1 and 3). XOR is famously a nonlinearly separable function, whereas AND or OR are linearly separable, implying nonlinear vs. linear transformations of the same inputs in the considered architectures (*21–23*) (see Figures 1A and 3). We aimed to reverse engineer the different stages of linear and nonlinear computations in brain networks that implement the algorithms (*24*). To do so, we simultaneously presented the inputs laterally within the visual field (a pair of sunglasses on a face, with dark ‘on’ vs. clear ‘off’ lenses representing the binary inputs, Figure 1A) so that occipital cortex initially represented each separately and contra-laterally (i.e. with left vs. right input projected in right vs. left occipital cortex; see *Methods, Stimuli*). Our analyses of the ensuing computation in the networks of individual participants systematically revealed four stages of computation that represent and transform the same inputs to produce different task-specific behavior (N = 10 participants per task, each analyzed separately to provide an independent replication; we further replicated the key results in different participants, using opposite-phase Gabor patches and also with sequentially presented inputs) (*25–28*).

**Figure 1.**
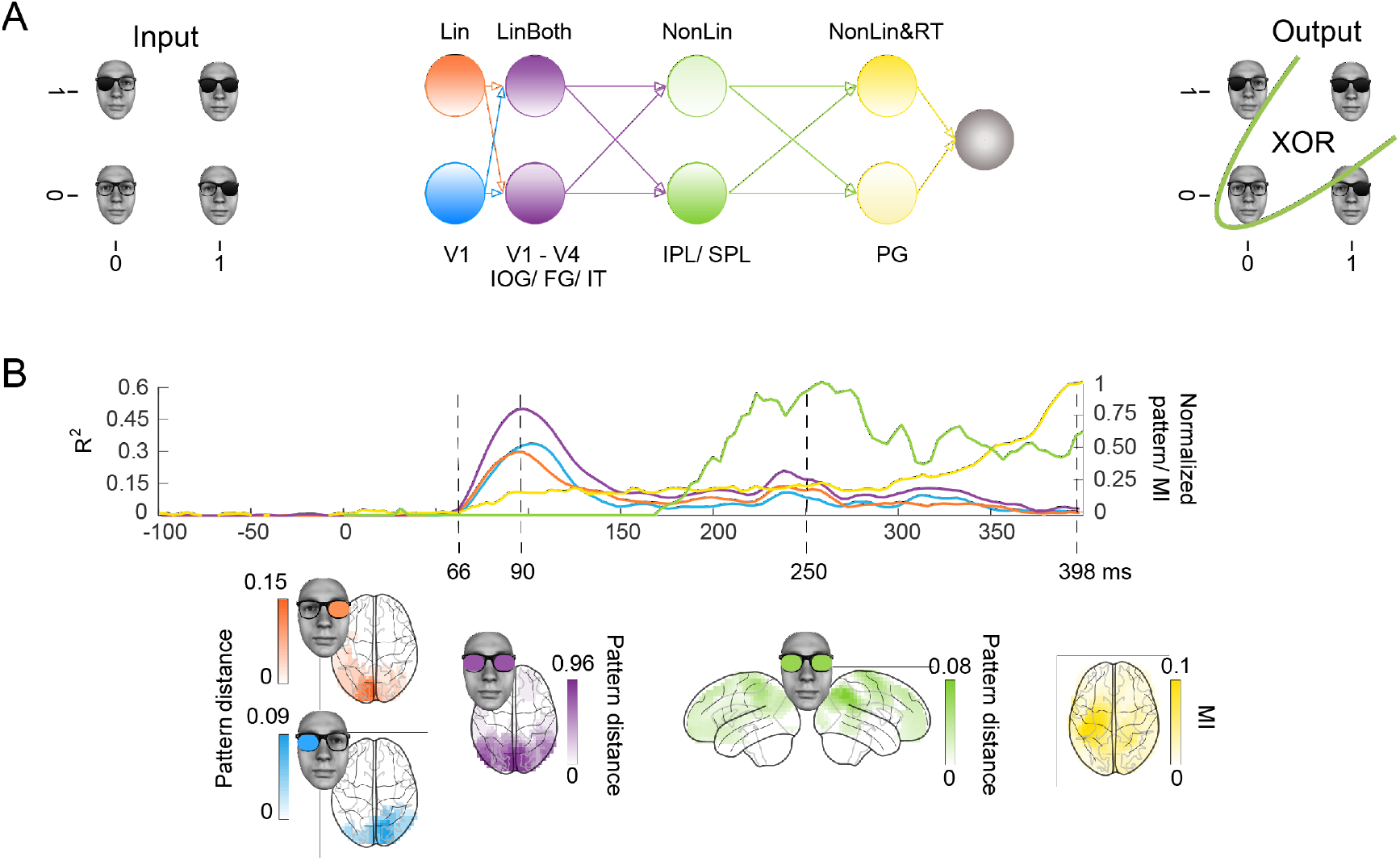
**(A) Hypotheses and schematic hierarchical brain network in the XOR task.** Stimuli consisted of the image of a face wearing glasses, with dark (“1”, ‘on’) and clear (“0”, ‘off’) left and right lenses serving as inputs, for a total of 4 input classes for XOR behavioral decisions. **(B) Four hierarchical stages of computations.** Each colored curve shows the average (N = 10 participants) time course of the maximum across sources that: (1) linearly discriminates in its MEG activity the ‘on’ vs. ‘off’ state of the left (Lin, blue) and right (Lin, orange) inputs (multivariate R^2^), (2) linearly discriminates both inputs (LinBoth, magenta) (multivariate R^2^ of joint stimulus model with no interaction), (3) nonlinearly integrates both inputs with the XOR task pattern (NonLin, green) (XOR pattern metric) and (4) nonlinearly integrates both inputs with the XOR task pattern (see Methods) and with amplitude variations that relate to RT (Mutual Information, MI(MEG; RT)). (yellow). Colored brains localize the regions where these computations start (onset times for left and right) or peak (peak latencies for both, XOR and RT) (p< 0.05 FWER-corrected with a permutation test, see *Methods, Linear vs. Nonlinear Representations; Representation Patterns*). See Table 2 and Supplementary Figure S1 for individual participant replications of each computation in the same brain regions and time windows.

**Figure 2.**
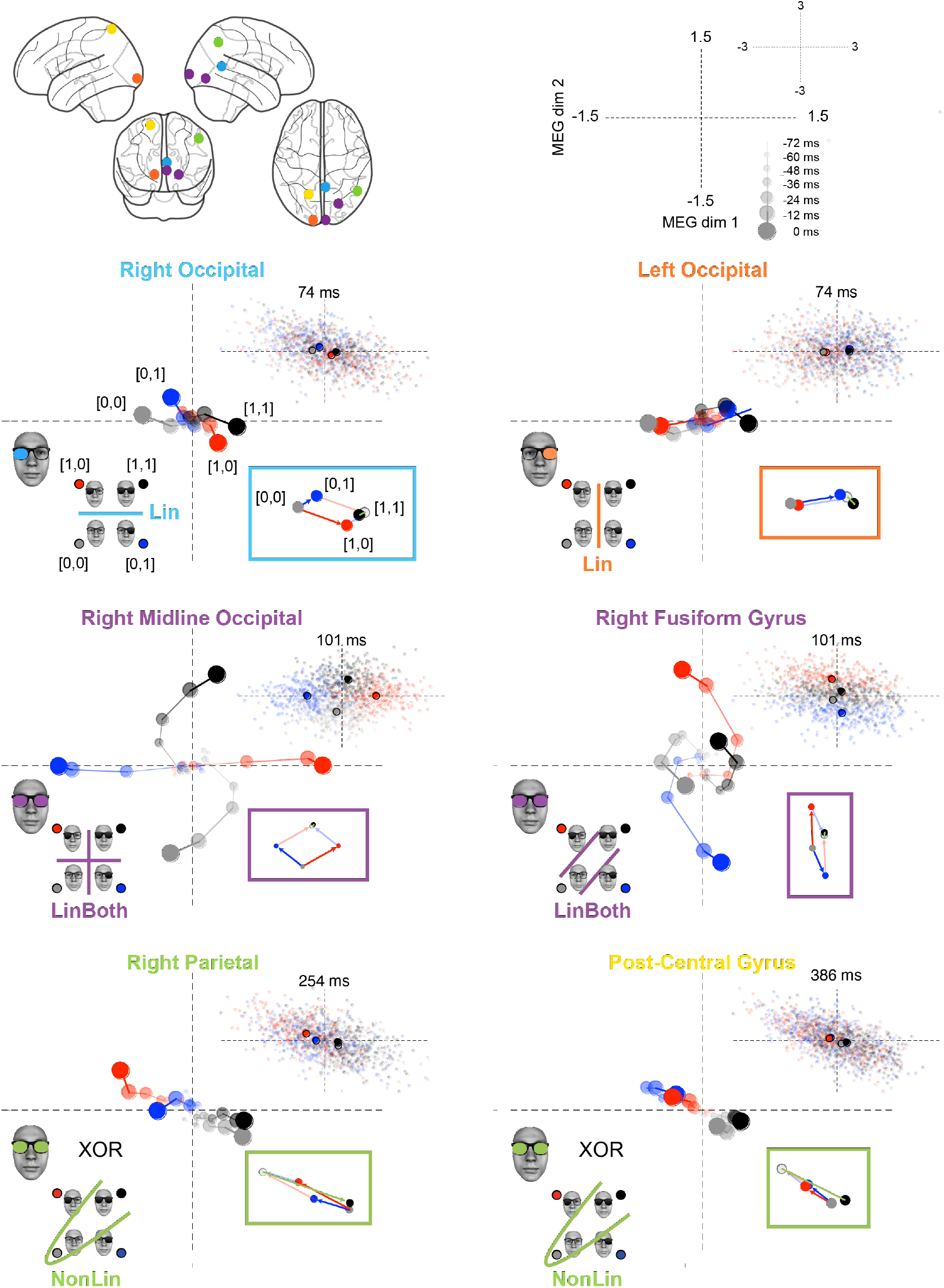
Dynamic unfolding of four different computations over different sources at different times from an example XOR participant. Each plot develops the computation of the color-coded source localized in the top glass brain. The axes represent the 2D source magnetic field response. Each small colored dot is single-trial source response. Larger dots show trial averages for each stimulus class, color-coded (see legend) and dynamically reported over seven timesteps corresponding to the seven triangular markers in Supplementary Figure S1A. Increasing timesteps of the dynamic trajectory is denoted with increasing dot sizes and saturations and connected with lines. In the legend, the color-coded discrimination line(s) vs. curve indicates respectively the linear (Lin, light blue and orange, BothLin, magenta) vs. nonlinear (NonLin, green and yellow) computations that the source represents at the final, seventh timestep (see adjacent scatter for the distribution of individual trials at this specific time). Inset vector diagrams on the right provide a geometric interpretation of the linear and nonlinear computations. Using the stimulus [0,0] as the origin: the blue arrow illustrates the source response to stimulus [0,1] (blue disk), the red arrow shows the source responses to [1,0] (red disk) and the grey disk illustrates the linear summation of these vectors (opaque lines). The black disc is the actual mean response to stimulus [1,1]. The green vector illustrates the discrepancy between the true response to [1,1], and the linear sum of the response to [0,1] and [1,0] responses and so illustrates directly the non-linear representation of stimulus [1,1].

**Figure 3.**
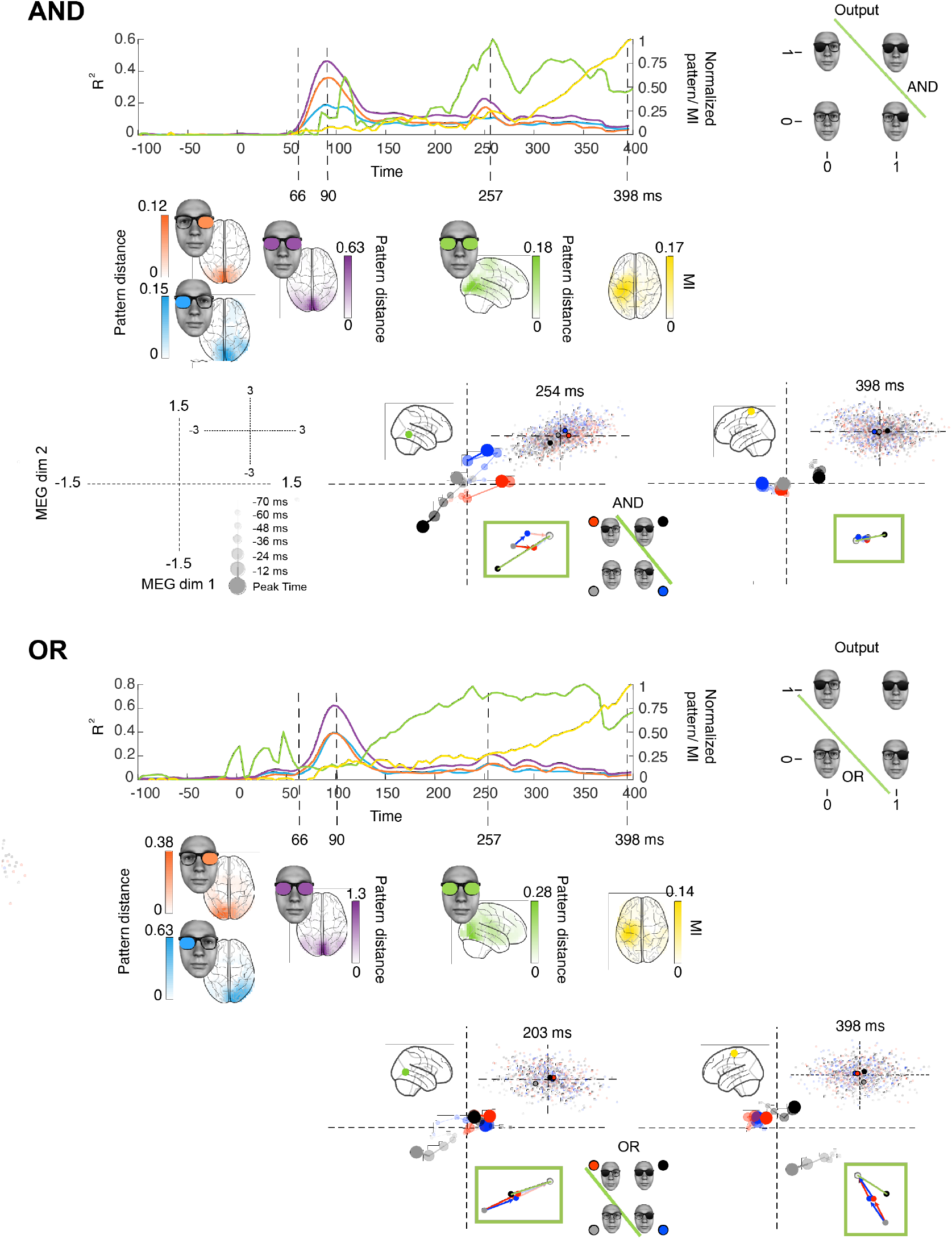
AND and OR tasks. Each colored curve averages (N = 10 participants) the time courses of the maximum across sources that: (1) linearly discriminates (R^2^) in its MEG activity the ‘on’ vs. ‘off’ state of the left (blue) and right (orange) inputs, (2) linearly discriminates both inputs (magenta), (3) nonlinearly integrates both inputs with the respective task pattern and (4) nonlinearly integrates both inputs with the respective task pattern and with amplitude variations that relates to RT (yellow). Colored brains localize the regions where these computations start (onset times for left and right) or peak (peak latencies for LinBoth, NonLin and RT) p<0.05 FWER-corrected with a permutation test, see *Methods, Linear vs. Nonlinear Representations; Representation Patterns*). See Table 2 and Supplementary Figure S1 for individual participant replication of each computation in the same brain regions and time windows. Single source plots in each task develops the data of one typical observer, where the green and yellow computations (cf. color-coded sources in the glass brain) differ between AND, OR and XOR (see Figure 2). Two-dimensional, source response to each input class is averaged, color-coded (see legend), and dynamically reported over the timesteps corresponding to the tick-marks of Supplementary Figure S1, AND and OR, and represented with varying dot sizes, saturations and connecting lines. In the legend, the color-coded discrimination curve indicates the nonlinear computations that source reflects at end time— i.e. the solution to AND and OR. NonLin computations at stages 3 and 4 were the only task differences between XOR, AND and OR.

## Results

Starting with behavior, Table 1 shows that participants were both accurate and fast in all tasks, with no significant task differences on average accuracy and reaction times (RT), measured with independent samples t-tests. We reconstructed the dynamic neural representation of the inputs of each participant from concurrently measured, source modelled magnetoencephalographic (MEG) activity (see *Methods, MEG Data Acquisition, Source Reconstruction*).

**Table 1.** Mean behavioral accuracy and median reaction times in XOR, AND and OR, 95% 95% percentile bootstrap confidence intervals in brackets. All pairwise comparisons, p > 0.05.

|  | ACCURACY | REACTION TIME |
| --- | --- | --- |
| **XOR** | 98.5% [97, 99] | 499 ms [461, 557] |
| **AND** | 99% [98.6, 99.5] | 457 ms [393, 505] |
| **OR** | 99.3% [99.2, 99.5] | 490 ms [439, 541] |

To simplify presentation, henceforth we use vector notation to denote the state of the two inputs and for example write left input ‘on,’ right input ‘off’ as [1,0]. To preview the analysis and key results, for each source and every 4 ms we fit linear models to explain the 2D MEG magnetic vector field activity in terms of the two presented binary inputs, with and without a non-linear interaction term between them. The interaction term captures the nonlinear integration of the two inputs on this MEG source and time point—i.e. when source response to [1,1] differs from the sum of the responses to [1,0] and [0,1]. Additional metrics quantified how the 2D MEG responses match the response pattern expected in each task (see *Methods, Representational Patterns*). Our analyses reveal that individual MEG source responses reflect changing representations of the visual inputs in the brain, revealing four different stages of neural computations that lead to behavior in each task (see *Methods, Linear vs. Nonlinear Representations, Representation Patterns*).

### Four systems-level stages of computation link stimulus to behavior

Figure 1B (XOR) and Figure 3 (AND and OR) show the time course of these four stages of computation averaged across the 10 participants of each task (Supplementary Figure S1 shows the results of each individual participant). Each task shows a similar dynamic unfolding: The first two stages represent and linearly discriminate the visual inputs; the third and fourth stages nonlinearly integrate them in a task-specific manner, revealing the solution of each task in the responses of individual MEG sources. The network model of Figures 1A schematizes these stages. Specifically, we show

1. **Linear, contra-lateral discrimination of each input state separately** (‘Lin’) in V1-4 regions with onset from ~60 ms post-stimulus (quantified with the multivariate R2 of a linear model for each binary input, color-coded in light blue for the left input, in orange for the right input, see *Methods, Linear Representation*).
2. **Linear discrimination of both inputs** (‘LinBoth’) on the same occipital and ventral sources ~100 ms post stimulus (quantified with the multivariate R2 of a linear model considering both inputs with no interaction, color-coded in magenta, see *Methods, Linear Representation*).
3. **Nonlinear integration of both inputs** (‘NonLin’) for task performance (XOR, AND or OR) in temporal-parietal regions ~260 ms (quantified with XOR representation pattern metric color-coded in green, also thresholded by significant improvement in model fit with interaction term, see *Methods, Nonlinear Representation*).
4. **Nonlinear integration of both inputs together with response-related activity** (‘NonLin&RT’) in post-central gyrus ~400 ms (quantified with Mutual Information between the 2D MEG magnetic field and reaction time on the corresponding trial, color-coded in yellow, also thresholded by XOR pattern metric and model interaction term, see *Methods, Information Theoretic Analyses*).

In Figures 1 and 3, colored sources shown in glass brain localize the regions where each color-coded computation onsets or peaks (cf. dashed lines) in different time windows post-stimulus. Table 2 shows independent replications of each computation within these regions and time windows, in at least 9/10 participants. We also replicated each color-coded computation at the level of individual participants, using Gabor inputs in XOR (Supplementary Figure S2) and a sequential presentation of the inputs in XOR, AND and OR (Supplementary Figure S3–4).

**Table 2.** Number of individual participant replications of the four color-coded computations, within the same region and time window: Lin, left and right occipital sources, [74-117 ms]; LinBoth, occipital mid-line and right fusiform gyrus, [74-117 ms]; NonLin, XOR: parietal, [261-273 ms], AND: temporal-parietal, [246-304 ms], OR: temporal-parietal [261-304 ms]; RT, post-central gyrus, [386-398 ms]. Bayesian population prevalence (*25*) of 9/10 = 0.89 [0.61 0.99]; 10/10 = 1 [0.75 1] (MAP [95% HPDI]).

|  | Lin | LinBoth | NonLin | NonLin & RT |
| --- | --- | --- | --- | --- |
| **XOR** | 10/10 | 10/10 | 9/10 (parietal) | 9/10 |
| **AND** | 10/10 | 10/10 | 10/10 | 9/10 |
| **OR** | 10/10 | 10/10 | 10/10 | 10/10 |

### Detailed dynamic unfolding of each computation at single source level

We next detail the dynamic unfolding of each color-coded computation on single sources, using an exemplar XOR participant (highlighted with colored curves in Supplementary Figure S1A, XOR). The selected sources maximize the metric of each computation—i.e. Lin: onset; LinBoth, NonLin, NonLin&RT: peak. The glass brain in Figure 2 locates the selected sources and color-codes them by type of computation. The panels of Figure 2 visualize the dynamic response trajectories of each source to the same four stimuli (representing the ‘on’ vs. ‘off’ combinatorics of the two inputs) over 72 ms, with a 12 ms timestep resolution (those indicated with triangle markers in Figure S1A). To preview the results, different source response trajectories to the same inputs detail the neural implementation of the different color-coded computations.

To illustrate, we start with the light blue right occipital source (top left panel). Its Lin computation develops over 7 time-steps (from 2 to 74 ms) to linearly discriminate the ‘on’ (dark) vs. ‘off’ (clear) state of the left input (see schematic in the bottom left quadrant). The plot shows how 2D source activity progressively separates left lens ‘on’ trials (red and grey stimuli, see blue line in schematic) from left lens ‘off’ trials (blue and black stimuli). The adjacent scatter (upper right quadrant) shows the source response to individual trials at 74 ms (final plotted time point). The vector diagram (lower right quadrant) confirms that the linear addition of the vector responses [0, 1] and [1, 0] (gray point, sum of blue and red vector) is close to the actual source response to [1, 1] (black point). The left occipital source (orange) reflects a similar unfolding for the linear discrimination of the right input ‘on’ vs. ‘off’ state. Again, the response is close to the linear sum of the responses to each individual input.

The second computation (LinBoth, magenta) that linearly and jointly represents the ‘on’ vs. ‘off’ state of both inputs takes two distinct forms. In midline occipital sources, all four stimuli are equally spread out in the quadrants of the source response space (i.e. all inputs are equally discriminated). In contrast, the right fusiform gyrus source discriminates the [1,0] and [0,1] stimuli with an opposite activity, whereas the [1,1] (black dot) and [0,0] (grey dot) stimuli are less discriminated. The vector diagrams of the two LinBoth examples confirms that the joint response to [1,1] is indeed the sum of [1,0] and [0,1] responses. Interestingly, the two LinBoth discriminations illustrate a progression towards an XOR representation. The first LinBoth midline occipital source discriminates equally each input in the quadrants of its 2D response. In contrast, the amplitude response of the LinBoth right fusiform gyrus source can linearly discriminate the XOR responses, but only when a nonlinear operation is added (i.e. drawing a circle that separate the two ‘same’ black and grey stimuli near the original in the 2D source response space from the two ‘different’ blue and red stimuli). So, the right fusiform gyrus LinBoth stage likely represents an important intermediate step towards the computation of XOR. We will see next that the green computation adds this nonlinear computation.

The third and most critical computation that starts distinguishing the task (NonLin, green) occurs when the parietal source nonlinearly represents the XOR solution for behavior, with ‘same’ vs. ‘different’ stimuli discriminated at 254 ms. Like LinBoth in right fusiform gyrus, this representation has black dot [1,1] and grey dot [0,0] responses close together, but with two critical differences. First, the red and blue vectors (lower right quadrant) now point in the same direction, rather than in opposite directions, as happens in right Fusiform Gyrus LinBoth. Such different source-level responses to the same [1,0] and [0,1] stimuli likely reflect different activities of the neural populations in the regions where the two sources are located. In parietal source NonLin, responses to [1,0] and [0,1] stimuli are magnetic fields with the same dipole orientation, suggesting that the same pattern of cortical activity (i.e. the same neural population) responds to both stimuli. In right fusiform gyrus source LinBoth, responses to [1,0] and [0,1] are magnetic fields with different dipole directions, suggesting that different neural populations within the region modelled by the source respond to each stimulus. Second, the representation of [1,1] (black dot) is now highly non-linear (green vector), lying far away from the sum of the red and blue vectors of the individual inputs. Following this nonlinear transformation, the XOR outputs are now linearly decodable in the 2D MEG response space.

Finally, the fourth stage on a post-central gyrus source (NonLin&RT, yellow) also nonlinearly represents the stimuli, also allowing linear readout of the XOR task outputs at 386 ms. In addition, this source activity now relates trial-by-trial to behavioral reaction times.

Figure 3 shows that the key differences in the AND and OR tasks are at the third and fourth stages, where the temporal-parietal (green) and post-central gyrus (yellow) sources represent AND and OR solutions for task behavior. The earlier stages linearly represent the two inputs, with light blue and orange Lin, magenta LinBoth discriminating the four stimuli as in XOR participants (see Supplementary Figure S1 for individual replications, prevalence = 1 [0.75 1]). In NonLin and NonLin&RT stages, the representation is non-linear and reflects the task (see vector diagrams inset bottom right quadrant). In AND, the task output is linearly separable in the 2D MEG response space: the black [1,1] response is further from the other stimulus classes than they are from each other, see *Methods, Representational Patterns*. Similarly in OR, the task outputs are linearly separable, with the grey [0,0] stimulus class represented apart from the other ones. This shows how these later post-200ms computational stages involve non-linear task-specific stimulus representations in ventral (NonLin) and parietal (NonLin, NonLin&RT) areas.

## Discussion

Here, we addressed the challenge of understanding where, when and how the brain dynamically implements different algorithmic computations over sensory inputs. We tightly controlled behavior using the simple logical functions XOR, OR and AND as tasks that require different computations over the same tightly controlled binary inputs. Our analyses revealed, at the level of individual MEG sources, four main stages of computation that dynamically unfold from ~60 ms to 400 ms post-stimulus. The first computation linearly discriminates the ‘on’ vs. ‘off’ state of each input in contral-lateral occipital cortex ~60 ms post-stimulus. This is followed by the linear discrimination of both inputs on occipital and ventral sources ~100 ms, followed by the nonlinear integration of both inputs, revealing the XOR, AND or OR task solution in 2D source response space in the parietal-temporal regions ~260 ms, and finally the nonlinear integration with RT-related activity in post-central gyrus ~400 ms. These four stages are common to XOR, AND and OR, with the main task-related changes occurring in the latter two non-linear stages. Notably, we performed all statistical analyses leading to these results within each individual participant, controlling the family-wise error rate over all considered sources and time points. By treating each participant as an N-of-1 study, 10 participants per task provide 10 independent replications of the experiment. We replicated the four computational stages in at least 9/10 participants (and in two further replication experiments with similarly high prevalence), providing strong evidence that a majority of individuals in the population sampled and tested in the same way would show the same effects (*25*).

### Reverse engineering systems-level algorithms

Our systems-level approach aims to reverse engineer, from mass brain signals, the hierarchy of brain computations that represent and transform sensory inputs to produce behavior—that is, the brain’s algorithm of the behavioral task. The four stages of computation that we systematically found in each individual participant and tasks meet the five key properties of an algorithm: (1) The inputs were specified as the four possible combinations of two binary inputs; (2) The output responses were also specified as the responses of the logical functions XOR, OR and AND; (3) The algorithms were definite in each task, with a sequence of two characterized Lin and two NonLin computations that transformed the same inputs into the task-specific outputs; The algorithms were also (4) effective in the brain, in the sense that they only relied on brain resources and (5) finite in processing time, producing behavior with ~450-500ms.

Note that reverse engineering a systems-level algorithm at the granularity of MEG brain measures does not preclude the existence of different compositions of algorithms at lower levels of granularity of individual neurons that together implement the higher level algorithm (much like the lower granularity algorithms of machine language implement the higher-level algorithms of C++). Rather, such systems-level analysis provides constraints on where (the brain regions) and when (the specific time windows) specific computations take place, enabling targeted studies of the algorithmic “how” across modalities and granularities of brain measures. Jonas and Kording (2017) used a related systems-level approach to understand the hierarchy of computations of a micro-processor and concluded that there was risk that analytic approaches in neuroimaging could fall short of producing a meaningful algorithmic understanding of neural activity. We could do so here because we adhered to the main properties of an algorithm: our explicit behavioral tasks (i.e. XOR, OR and AND) require an implementation of a specific computation on simple (i.e. fully characterized) binary inputs to achieve behavior. We could therefore trace the dynamic representations of the inputs into the 2D space of MEG activity to understand the stages of representation underlying the computation (i.e. Lin, LinBoth and task-specific NonLin). Such descriptive models of an algorithm enable explicit testing of the different stages of the computation hierarchy. For example, by manipulating the timing or nature of the presented stimuli, or by targeting causal interventions (e.g. magnetic stimulation, or stimulus manipulations) at specific brain regions and peri-stimulus times.

### Generalization to naturalistic categorization tasks

Generalizing from our case study of the algorithms underlying the simple XOR, AND and OR functions to more naturalistic face, object and scene categorization tasks will incur many challenges that we can frame in the context of the properties of an algorithm detailed above.

A key challenge is that the task-relevant features of real-world faces, objects and scenes, may be completely different for different behaviours and participants, effectively changing the inputs to the algorithm. Unfortunately, task- or participant-specific features are generally not considered in studies of neural representation, processing and categorization. Their understanding remains a similar challenge for Deep Convolutional Neural Network research, including instances when these are used as models of the brain. Specifically, a key property of an algorithm is that we specify its inputs as precisely as possible. In real-world categorizations, this implies understanding which specific features of the complex images are task-relevant for each particular participant performing each specific behavioral task. Furthermore, specification of the outputs is another key property of an algorithm. Passive viewing, or one-back tasks do not provide this specification. For example, from the same face, the feature of a smiling mouth feature will be used to overtly respond “happy,” but the forehead wrinkles to respond “45 years of age;” from the same car picture, its specific shape to respond “New Beetle,” but the front tyre shape to respond “flat tyre;” or the specific roof tiles to respond “Chrysler building” but the density of buildings on the horizon to respond “city” and so forth. Relatedly, experts vs. novices will use different features to classify the same pictures of the 35 different subtypes of sparrows that exist in North America. Such relative perceptual expertise and associated features generally characterize the relationship between visual cognition and outside world stimuli. Then, to infer the hierarchical stages of computation from the brain measures, we can start tracing the dynamic representation of these specific task-relevant input features, when we have formally characterized them, between stimulus onset and explicit output task-behavior, as we did here. Different modalities or granularities of brain measures (e.g. M/EEG, 3/7T fMRI, NIRS vs. single electrodes (*23*) and electrode arrays) will likely provide complementary understandings (e.g. timing vs. location) of the computations in different brain regions. And when we finally have a model of the computation hierarchy (even initially a descriptive model), we can test its key properties.

To conclude, we reverse engineered four stages of dynamic algorithmic computations over the same sensory inputs that produce different behaviors in the brain. We could do so because we explicitly controlled the input features and the explicit tasks that each individual participant was instructed to resolve while we modelled their brain response with MEG source level activity. Therefore, our results and methodology pave the way to study algorithmic computations when the stimuli and tasks are more complex (e.g. face, object and scene and their explicit categorizations) but well controlled (e.g. with generative systems rather than uncontrolled 2D images), as they are in any algorithm.

## Methods

### Participants

We recruited 35 participants (all right-handed; 24 females). All reported normal or corrected-to-normal vision and gave written informed consent. We conducted the study according to the British Psychological Society ethics guidelines and was approved by the ethics committee at the College of Medical, Veterinary and Life Sciences, University of Glasgow.

### Stimuli

We synthesized an average face using a generative photo-realistic 3d face model (*30, 31*) to which we added glasses with an image editing program (Adobe Photoshop). Black and clear lenses produced four different input conditions corresponding to four classes of logical inputs: 1) both clear, in vector notation [0,0], 2) left clear/right dark, [0,1]; 3) left dark/right clear [1,0]; 3) both dark, [1,1]. The edges of the left and the right lens were 0.5 deg of visual angle away from the centrally presented fixation cross.

### Task Procedure

Each trial began with a central fixation cross displayed for a randomly chosen duration (between [500-1000 ms]), immediately followed by one of the four stimulus classes described above and displayed for 150 ms. We instructed participants to maintain fixation on each trial, to pay attention to the left and the right lenses and to respond as quickly and accurately as possible by pressing one of two keys ascribed to each response choice with the index or middle fingers of their right hand.

Responses were “same” vs. “different” in the XOR task; “both dark” vs. “otherwise” in AND; or “at least one dark” vs. “otherwise” in OR.

Stimuli were presented in blocks of 80 trials, with random inter-trial interval of [800-1300 ms], and randomized stimulus order in each block. Participants completed a total of 20-24 blocks split across 2-3 single day sessions, with short breaks between blocks. Each session lasted 2.5-3 hours.

### MEG Data Acquisition and Pre-processing

We recorded the participants’ MEG activity using a 248-magnetometer whole-head system (MAGNES 3600 WH, 4-D Neuroimaing) at a 1017 Hz sampling rate. We discarded each participant’s runs with more than 0.6 cm head movement measured with pre- vs. post-run head position recordings. Participants were excluded if the number of trials remaining after preprocessing (artifact rejection and rejecting runs for excessive head motion) was less than 700. We excluded five participants resulting in a final sample sizes of N = 30 (10 per task). Mean head movement (averaged across blocks) across participants was 0.3 cm (min = 0.12, max = 0.44).

We performed analyses with Fieldtrip (*32*) and in-house MATLAB code, according to recommended guidelines (*33*). We high-pass filtered the data at 0.5 Hz (5^th^ order two-pass Butterworth IIR filter), filtered for line noise (notch filter in frequency space) and de-noised via a PCA projection of the reference channels. We identified noisy channels, jumps, muscle and other signal artifacts using a combination of automated techniques and visual inspection. The median number of trials for subsequent analyses was 1,064 (min = 701, max = 1,361).

Next, we epoched the data into trial windows ([−500 to 1000 ms] around stimulus onset, low-pass filtered the data at 45 Hz (3^rd^ order two-pass Butterworth IIR filter), resampled to 256 Hz, and decomposed using ICA, separately for each participant. We identified and projected out of the data the ICA sources corresponding to heartbeat and eye blinks or movements (2 – 4 components per participant).

### Source reconstruction

For each participant, we co-registered their structural MRI scan with their head shape recorded on the first session and warped to standardized MNI coordinate space. Using brain surfaces segmented from individual warped MRI, we then prepared a realistic single-shell head model. Next, we low-pass filtered the clean dataset at 40 Hz, re-epoched the data between −100 and 400 ms around stimulus onset, demeaned using a pre-stimulus baseline, and computed covariance across the entire epoch. Using average sensor positions across good runs (i.e. where head movement was <0.6 cm, see above), and a 6 mm uniform grid warped to standardized MNI space, we then computed the forward model, keeping the two orientations of MEG activity. We computed the Linearly Constrained Minimum Variance (LCMV) beamformer (*34*) solution with parameters “lambda = 6%” and “fixedori = no”. The resulting inverse filter applied to the sensor-space MEG activity enabled reconstruction of the single-trial 2D MEG magnetic field vector (i.e. dipole with amplitude and direction) activity time courses on 12,773 grid points. Using a Talaraich-Daemon atlas, we excluded all cerebellar and non-cortical sources, and performed the statistical analyses on the remaining 5,848 cortical grid points.

### Linear vs. Nonlinear Representations

#### Linear Representation

Every 4 ms time point between −100 and 400 ms post-stimulus, we computed independent multivariate linear regressions to model the dynamic representation of the state of each input (i.e. 0, clear lens, vs. 1, dark lens) into the 2D MEG responses of each source. We computed three linear models covering each input separately (Left, L, and right. R) and additively.

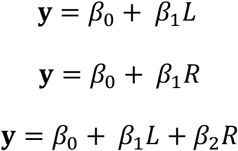

We fitted each model with ordinary least-squares, resulting in beta coefficients for the intercept and slope. We quantified the fit in the 2D response space of the voxel with a multivariate R^2^ that quantifies multivariate variance as the determinant of the covariance matrix:

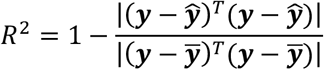

where ***y***,***ȳ***,***ŷ*** are the 2D source data, their mean and model predictions respectively. This linear modelling produced a time course of R^2^ values per source with 4ms resolution.

To control the Familywise Error Rate (FWER) over all considered time points and sources, we computed a non-parametric statistical threshold with the method of maximum statistic (*35*). Specifically, on each of 100 permutations we randomly shuffled input state (‘on’ vs. ‘off’) across the experimental trials, repeated the linear modelling and R^2^ computation explained above, and extracted the maximum R^2^ across all sources and time points. This produced a distribution of 100 maximum R^2^ values, of which we used the 95^th^ percentile as statistical threshold (FWER p < 0.05).

#### Nonlinear Representation

A fourth model considered, for each source and time point the nonlinear interaction term between the Left and Right inputs.

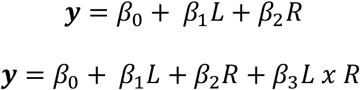

A log-likelihood ratio (LLR) tested whether the added interaction term significantly improved model fit (p < 0.05, FDR-corrected over time points and sources, (*35, 36*).

#### Representation Patterns

Linear and nonlinear representations of the two inputs into 2D voxel activity could form a variety of different patterns. To ensure that these patterns corresponded to expectations (e.g. of an XOR solution), we applied two further computations at each source and time point. First, we computed the pairwise Mahalanobis distances as detailed below between the color-coded 2D distributions of single trial MEG activity in response to each input class (see Figure 2). To do so, we averaged the covariance matrices of each pair of input conditions and multiplied the inverse average covariance by the difference of the condition means:

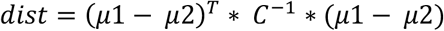

Then, we quantified the geometric relationships between the two-dimensional centroids of the source responses to each input class. We did so by combining the pairwise distances in the way that quantifies the expected representational pattern (see Figure 5, right):

- left lens representation (LL): mean([d1, d3, d5, d6]) – mean([d2, d4]). This measure contrasts distances where the left lens state changes, with distances where the left lens state does not change.
- right lens representation (RL): mean([d1, d2, d4, d6]) – mean([d3, d5]). As above, for the right lens.
- both lenses representation (BL): mean(all) – std(all)
- XOR representation: mean([d2, d3, d4, d5]) – mean([d1, d6]). Contrasts the distances between elements of the two different output classes with the distances between elements within each output class.
- AND representation: mean([d4, d3, d6]) – mean([d1, d2, d5])
- OR representation: mean([d6, d2, d5]) – mean([d4, d3, d1])

**Figure 5.**
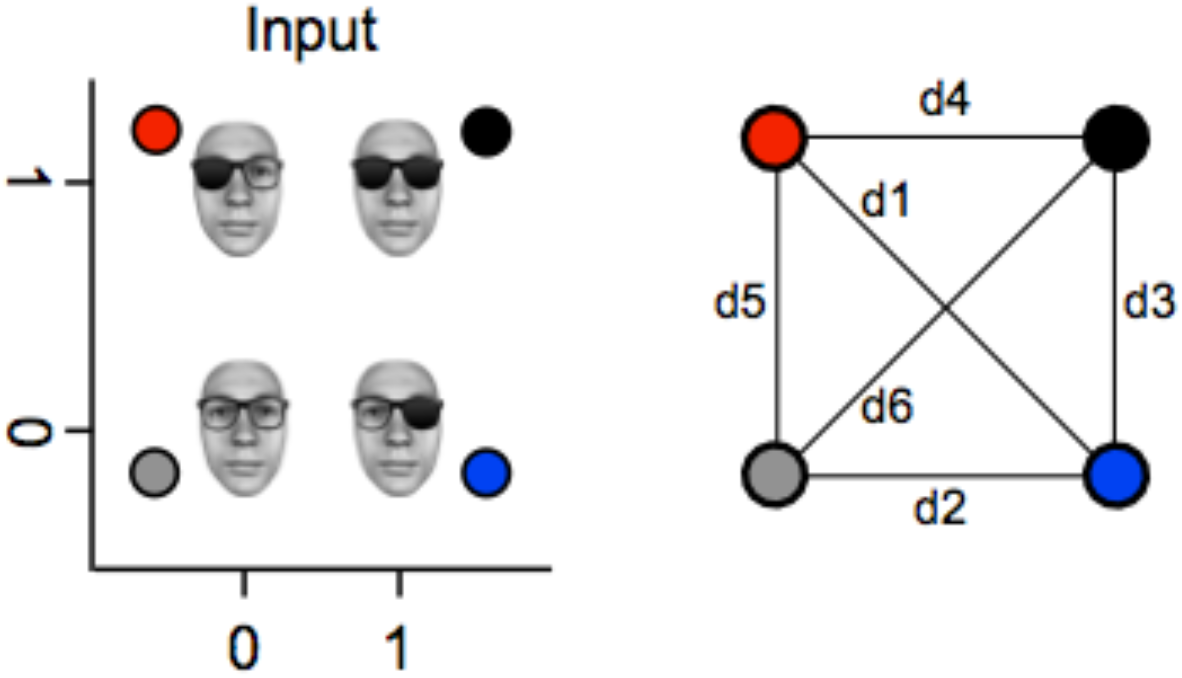
Note: the face stimulus was artificially synthesized and so does not belong to any real person.

We tested the statistical significance of each pattern with permutation testing, using again the method of maximum statistics. Over 25 permutations, at each source and time point, we randomly shuffled input ‘on’ and ‘off,’ repeated the above distance calculations, computed the maximum differences and used the 95^th^ percentiles of these maxima as thresholds (FWER p < 0.05, corrected).

Finally, we applied the conjunctive thresholding of significant R^2^ for linear representation patterns of the left or right input (see *Methods, Linear Representation*) with significant LL, RL and BL pattern (see above), and the conjunctive thresholding of significant nonlinear LLR test statistic (see *Methods, Nonlinear Representation*) with significant XOR, AND, and OR representation patterns (see above).

#### Localization of Representation Patterns

We quantified the temporal dynamics of the four stages of information processing in individual participants as follows. For early representation of the left and right lenses, we computed representation onsets as the first significant time point of R^2^. For early representation of both lenses, we computed its R^2^ peak time. Finally, for XOR, AND and OR nonlinear representations, we also selected the peak times of the respective representation task pattern distance measure. For each state, we then computed time stamps as median across observers and extracted the sourcewise representation averaged across participants at the corresponding time stamp. We plotted these sources back onto glass brains in Figure 1C using *Nilearn* (*37*).

#### Information Theoretic Analyses

We used information theory to quantify the association between RTs and MEG activity, as MI<RT; MEG>, splitting RTs into 4 equiprobable bins and using continuous MEG (on all sources and time points). To this end, we used Gaussian-Copula Mutual Information (GCMI; (*38*) on all voxels and time points. We assessed statistical significance with a permutation test.

## Acknowledgments

P.G.S. received support from the Wellcome Trust (Senior Investigator Award, UK; 107802) and the Multidisciplinary University Research Initiative/Engineering and Physical Sciences Research Council (USA, UK; 172046-01). R.A.A.I. was supported by the Wellcome Trust [214120/Z/18/Z]. The funders had no role in study design, data collection and analysis, decision to publish or preparation of the manuscript.

## Supplementary Results

**Supplementary Figure S1.**
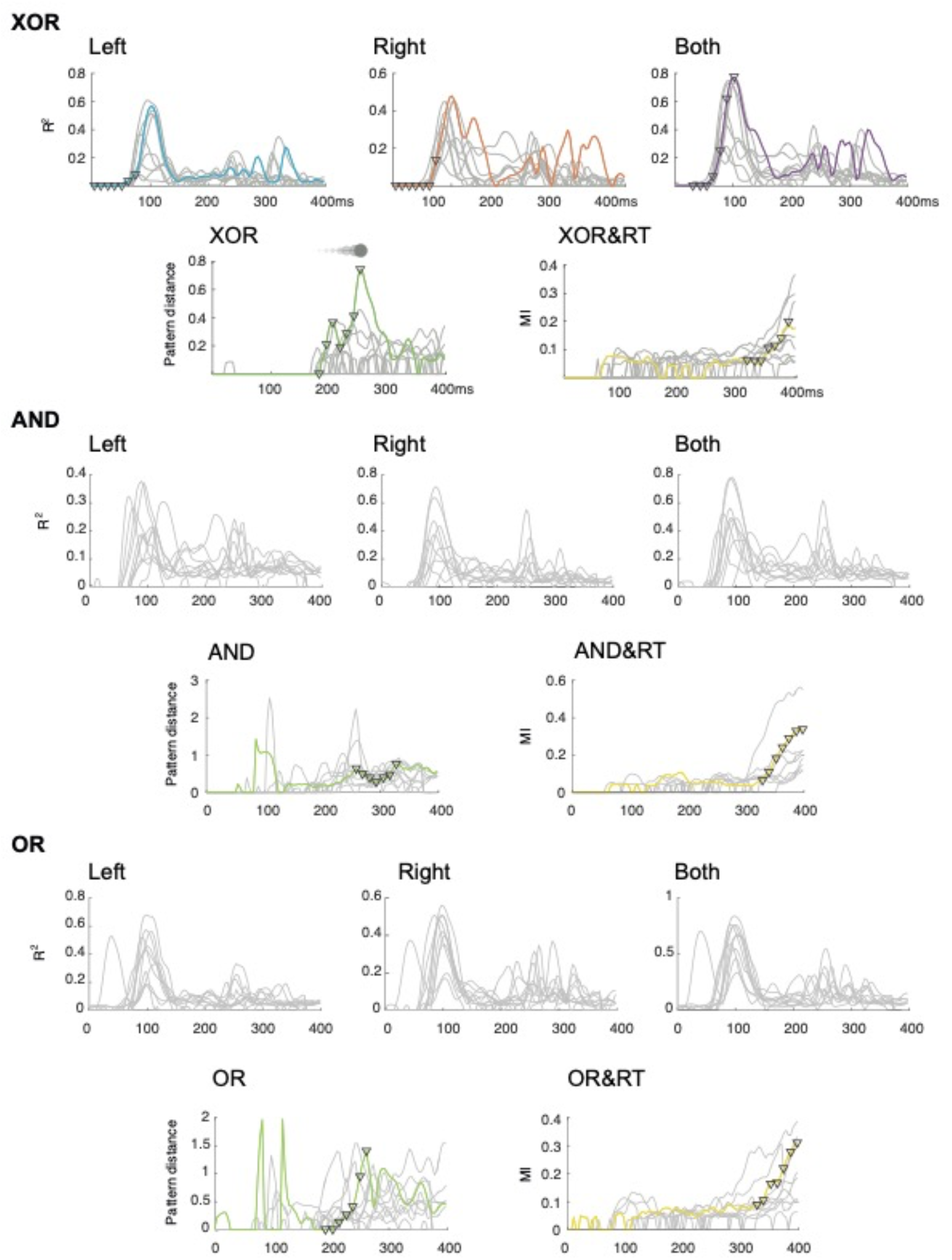
Results for individual participants. Grey curves correspond to the individual participants’ timecourses of the individual source Lin (left and right), LinBoth, NonLin and NonLin and RT, in the XOR, AND and OR tasks. Only significant (FWER p=0.05) time points are shown, other time points are set to 0. The colored lines correspond to the data of one exemplar participant used in Figure 2 (XOR) and 3 (AND and OR). Tick marks on each colored line correspond to the seven time steps of the dynamic unfolding of each computation.

**Supplementary Figure S2.**
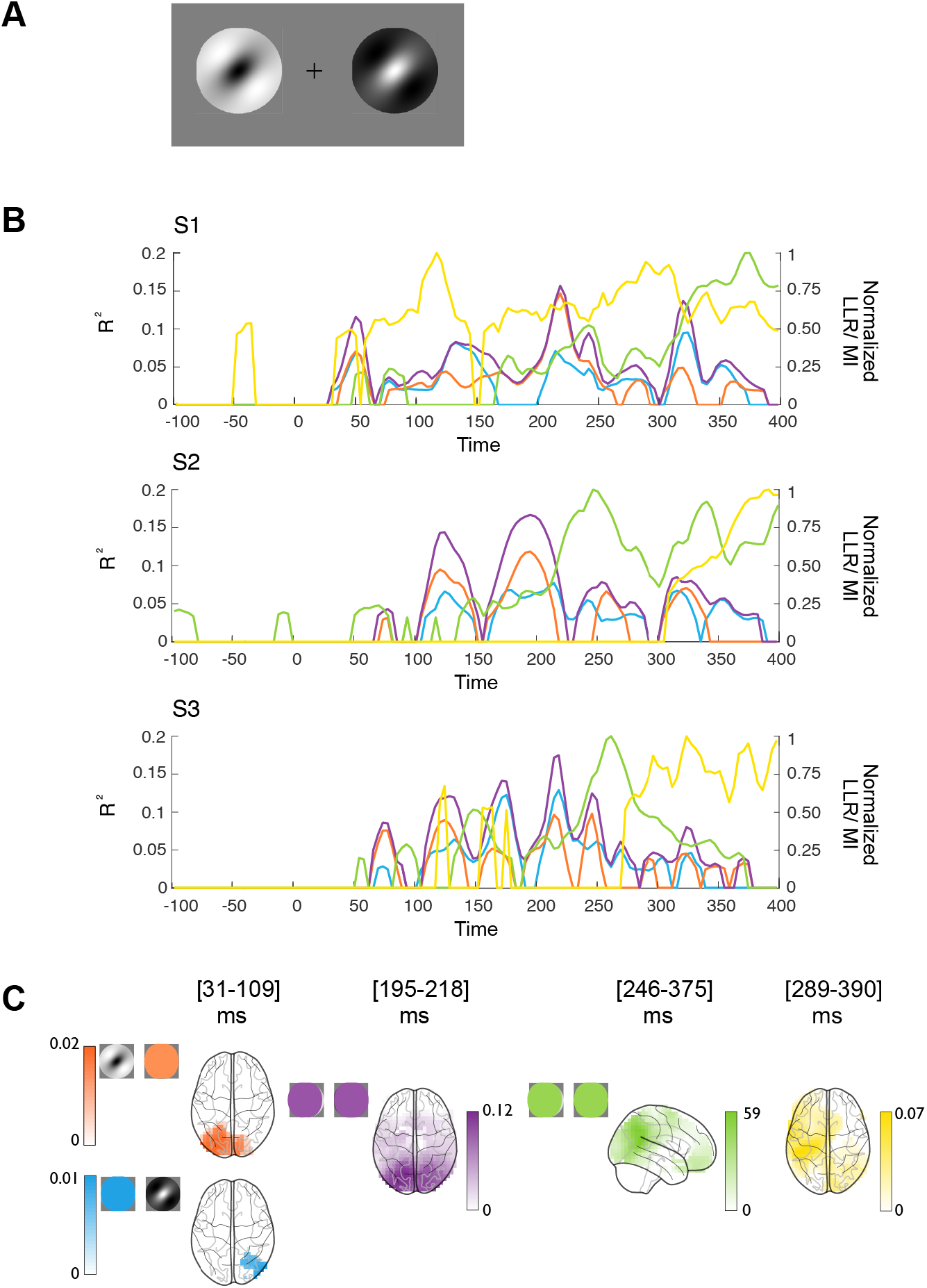
Replication of four stages of computation with Gabor stimuli. **(A) Stimuli.** In three participants, we presented two spatially separated Gabor and phase-opposed Gabor patches. Participants responded when the two patches had the same vs. different color with different keyboard keys (an XOR task). Gabors were presented in four randomly chose orientations, spanning in total 11 × 11 deg of visual angle (the inner edges of each Gabor were located ~5.7 deg of visual angle away from the centrally presented fixation cross). Gabor inputs were simultaneously presented on the screen for 150 ms. **(B) Four hierarchical stages of computations.** Each plot shows the maximum within participant of source-level (1) linear discrimination of each Gabor input Lin, (blue and orange), (2) linear discrimination of both input, Linboth (magenta), (3) nonlinear integration of both inputs, NonLin (green) with the XOR task pattern and (4) with amplitude variations that relate to RT NonLin&RT (yellow). **(C) Colored brains** localize the regions where these computations start. The brackets report the per participant onset times (for Lin, left and right) and peak latencies (for LinBoth, NonLin XOR and XOR&RT), and averaged across participants in the colored brains.

**Supplementary Figure S3.**
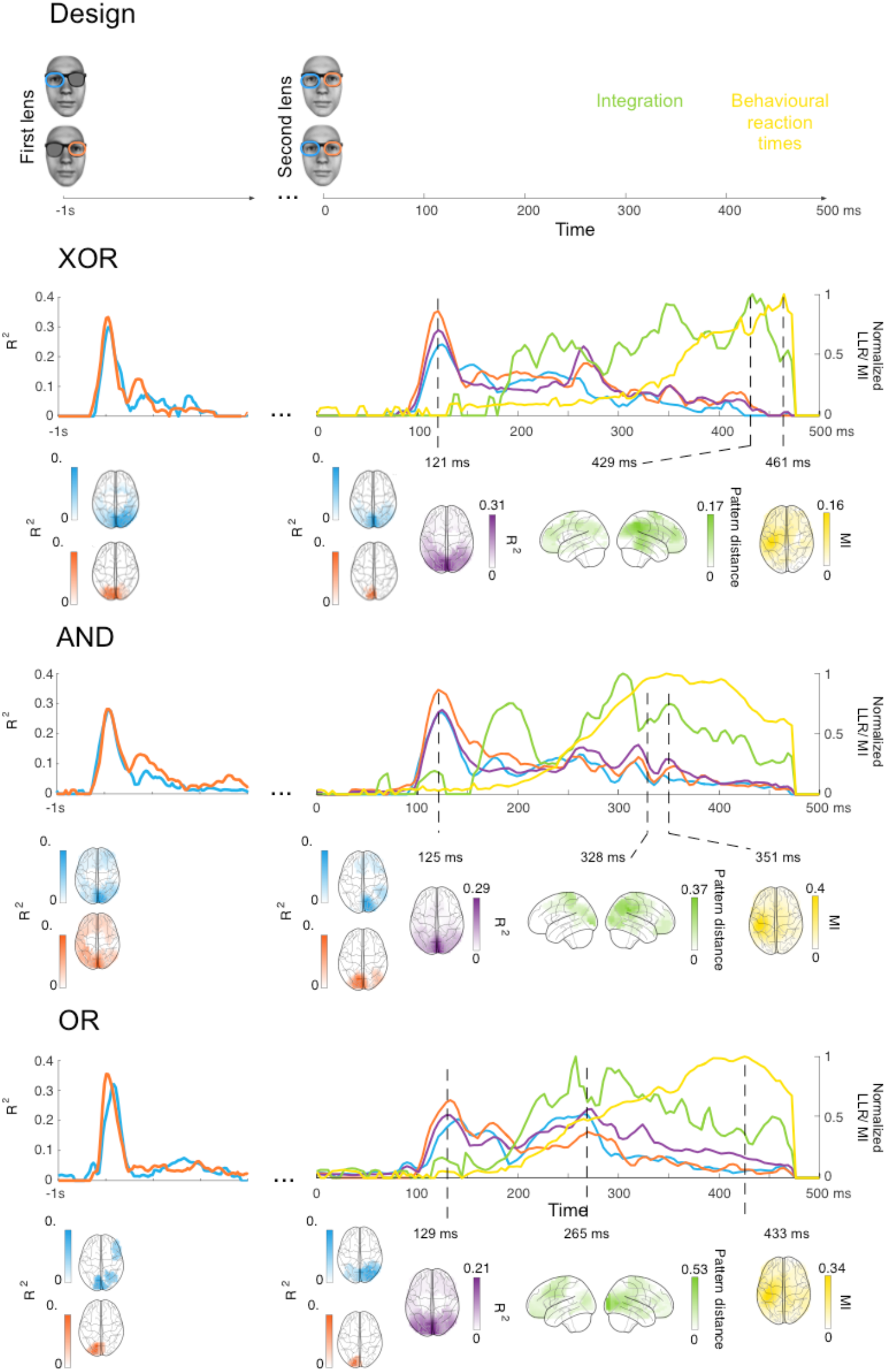
Four stages of computations replicated with delayed stimulus presentation. Using the same face stimuli as in the main experiment, we presented either the right or the left input as grey for the first 1000 ms following stimulus onset. The other input was either clear or dark. After 1000 ms, the grey input changed into either clear or dark. Participants waited until the grey input changed, and then responded according to the logic-gate task: XOR (N=5, top), AND (N=4, middle), and OR (N=5, bottom) (2 participants were excluded under the same criteria as the main experiment). Linear representation of the first input started around 80 ms post-stimulus (with a ~100 ms peak; or around −900 ms with respect to second lens presentation). The second input was linearly represented from ~100 ms with a peak at ~120 ms (blue, orange and magenta time courses). Averaged time courses of the Lin and NonLin computations across individual participants (see Supplementary Figure S4) replicated the results of Figure 1.

**Supplementary Figure S4.**
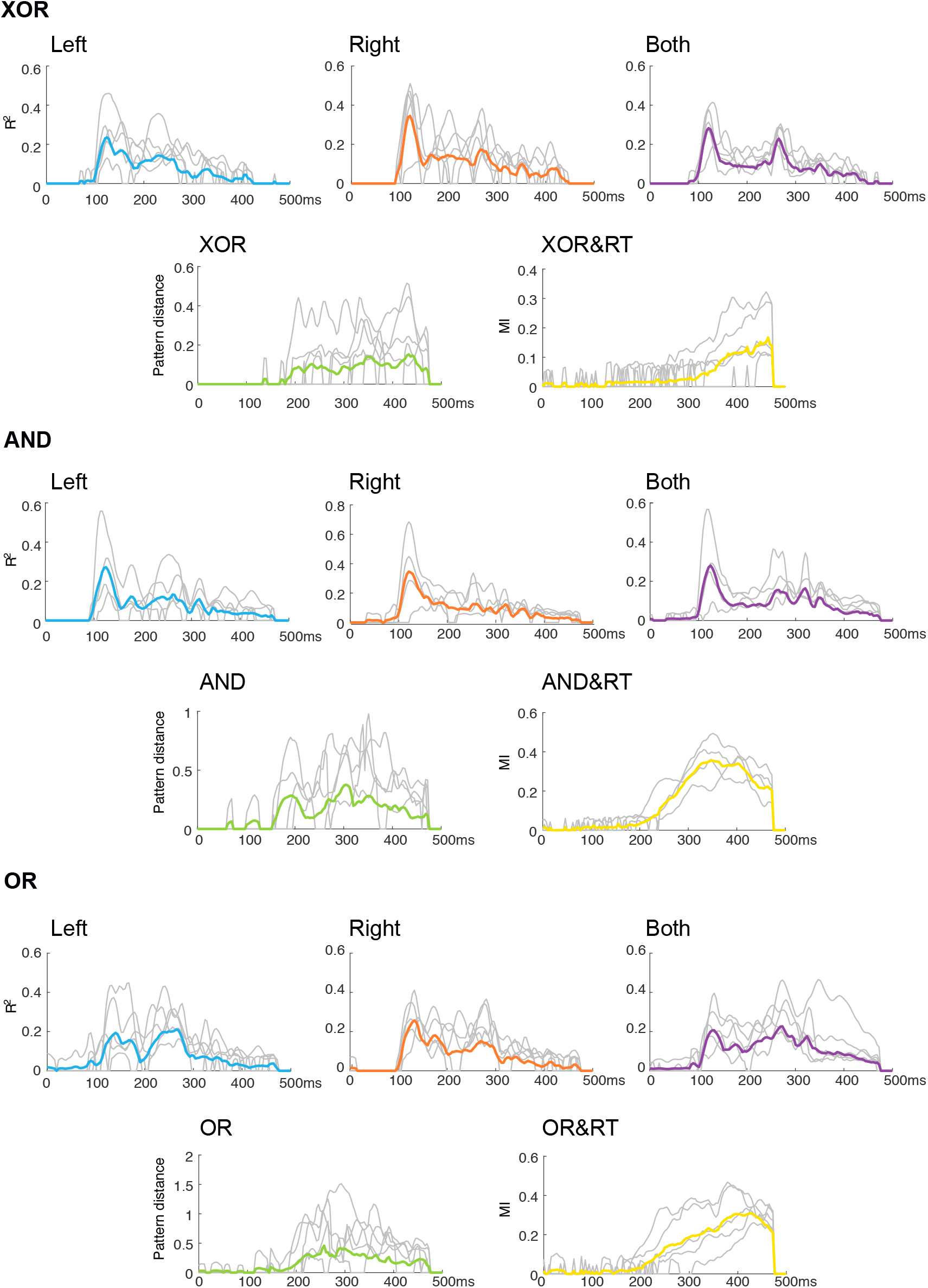
Four stages of computations replicated with delayed stimulus presentation: Individual participants results. Grey curves correspond to the individual participants’ timecourses of the individual source Lin (left and right), LinBoth, NonLin and NonLin&RT, in the XOR, AND and OR tasks, with 0ms corresponding to the time point when the second input is revealed (i.e. when the grey lens turns clear or dark, cf. Supplementarty Figure S3). The colored lines correspond to the mean across participants.

